# Identifying priority connectivity in a tropical forest hotspot severely affected by land use changes

**DOI:** 10.1101/2020.03.13.991372

**Authors:** Neil Damas de Oliveira-Junior, Gustavo Heringer, Marcelo Leandro Bueno, Vanessa Pontara, João Augusto Alves Meira-Neto

## Abstract

The probability that a propagule reaches a certain location where it can establish and persist is affected by the distance from the source. Fragmented landscapes often promote the isolation of habitats that hinder or impede the movement of species that affect their range distribution, even when environmental conditions are adequate. Studies that assess the connectivity of the landscape are essential to ensure that ecological conservation planning covers all vital processes that occur on a landscape scale. Based on the classification of land cover in the Rio Doce Basin - RDB, we used a habitat / non-habitat approach to assess RDB connectivity for tree species. Based on circuit theory, we built 6 surface models of resistance based on habitat and non-habitat areas. We performed the analysis using the GIS plug-in and Linkage Mapper to generate lest cost path maps. The RDB is very fragmented, but still has functionally connected regions. The west to nothwestern and southeastern portions of the basin are well-connected and demand conservation practices, while the center-north of the basin and the far southwest are regions with greater resistance to connectivity as a result of strong anthropic pressures that reduce forests, requiring intervention through restoration projects avoiding loss due to lack of connectivity. In addition, land reclamation projects in degraded areas must also be organized in the Linhares region, as it is inserted in a region of very high biodiversity with a high level of endemism and a high number of threatened species.

## Introduction

Connectivity is a major concern in landscape ecology and land conservation (Saura et al., 2014). A well-connected vegetation patch receives more dispersers and also provides more propagules (Taylor et al., 1993). In addition, connectivity allows species moving in the landscape to collect resources, colonize new habitats and help maintain genetic diversity through the dispersion of pollen and seeds (Tumas et al., 2018). These ecological processes can be affected by the lack of connectivity, risking the permanence of plant populations in a landscape (Santos et al., 2019).

In a scenario of global change, connectivity is very important to alleviate the consequences of changes in environmental conditions, allowing species to move, change their distributions (Opdam and Wascher, 2004), and maintain biodiversity in fragmented landscapes (Matos et al. 2016). Therefore, studies assessing landscape connectivity are essential to ensure that ecological conservation planning covers all vital processes that occur on a landscape scale.

The probability that a propagule reaches a certain place where it can establish and persist is affected by the distance from a source. In other words, the greater the distance, the less likely a species is to reach an appropriate habitat and establish (Levey et al., 2008; Weber et al., 2014). Consequently, fragmented landscapes, such as the Atlantic Forest, promote the isolation of habitats that can hinder or prevent the movement of species, which affects their distribution and harms them when changes cause inappropriate conditions for the existence of those species (Magnago et al., 2015; Matos et al., 2017). In contrast, areas with adequate habitat, such as small patches of forest between larger fragments, act as corridors of species allowing their movement in the landscape (i.e. connectivity; e.g., Matos et al., 2019).

Modeling connectivity for plant species in a landscape is a challenge as there are many factors that affect functional connectivity in plants. The dispersal of pollen and seeds depends on biotic and abiotic factors, which in turn are affected by the landscape (Auffret et al., 2017). The resistance distance method solves part of these problems because it considers different levels of suitability for the dispersion of plants and animals (McRae and Kavanagh, 2006), from hostile environments to different patch mosaics (Thiele et al., 2018a). This approach is advantageous because it explains the random movements in the landscape, considering more than one possible path (McRae and Kavanagh, 2006; Thiele et al., 2018a), and providing solid data for conservation planning and decision making (Correa Ayram et al., 2014; Fuller et al., 2006).

Here, we used landscape connectivity to analyze the Rio Doce Basin (RDB), assessing the connectivity between fragments on the landscape scale. We used a comprehensive data set on the composition of tree communities across the RDB and based on the land cover classification. We also used a habitat / non-habitat approach to assess RDB connectivity for tree species, where habitat was defined as all native fragments and non-habitat was defined as all other types of land cover. This study aimed to identify areas whose lack of connectivity in the RDB needs attention for conservation planning in a biodiversity hotspot of tropical forests. Our general objective was to generate maps of the current fragments of native forest in the RDB. Finally, we used ground cover to generate maps showing areas with high resistance and showing the best fit for lower cost corridors (i.e., low resistance) using the Jaccard index of species composition as a proxy for connectivity.

## Materials and Methods

### Study area

The RDB, which occupies parts of the states of Minas Gerais and Espírito Santo, with a total area of 86,715 km^2^, between latitudes 17°45’ and 21°15’ S and longitudes 39°30’ and 43°45’ W. It is an important hydrographic basin where many important economic activities are developed, such as agriculture, livestock and mining. These activities are the main cause of fragmentation in this highly diverse region (Saiter et al., 2015). The Basin is comprised between Atlantic Ocean and Espinhaço Range from east to west, Negra and Aimorés Mountain in the north limits and Caparaó Range on southeast limits. In the RDB, the soils are predominantly Red-Yellow Latosols, and Red Cambisols being the first characterized by deep soils dystrophic with high saturation of aluminium and the latter very similar soil, but with variable depth (Marangon et al., 2013; Nunes et al., 2000; Soil Department of Universidade Federal de Vicosa, 2020). Most of the hydrographic basin is within the domain of the Atlantic Forest, with different physiognomies: tropical rainforest with well-distributed rainfall throughout the year and average temperature of 25°C, semi-deciduous tropical forest characterized by variable seasonality (Veloso et al., 1991), with dry season and rainy season with intense rains, mountains with rocky vegetation, mainly in the hydrographic basins. During the dry season, the average rainfall is 150 to 250 mm, and in the rainy season it varies between 800 and 1300 mm (Alvares et al., 2013).

The RBD is an important hydrographic basin where many important economic activities are developed, such as agriculture, livestock production and mining. These activities are the main cause of fragmentation in this highly diverse region, as well as the main cause of environmental disasters (Meira et al., 2016; Nazareno and Vitule, 2016; Saiter et al., 2015).

### Floristic data set

The tree occurrence matrix for 78 locations in the RDB (Fig. 1) was extracted from NeotropTree, a database containing a checklist of tree species in the Neotropics (Oliveira-Filho, 2014) and used as central areas in the analysis of connectivity (see below). The checklists were obtained from records of the occurrence of different sources of trees [a] Published floristic and quantitative research. [b] taxonomic monographs and [c] herbarium records available in the Flora and Fungi virtual herbarium (“speciesLink Network - Herbário Virtual da Flora e dos Fungos,” 2020) (INCT; http://inct.splink.org.br/). Information reliability, expert opinion and taxonomic literature were used to verify the data. Due to the high density of floristic and quantitative surveys for certain locations, the data was compiled and merged into a single checklist, being kept on separate checklists in locations where the vegetation does not remain constant within a radius of 5 km. As a result, a total of 1944 species distributed in 100 families, for a total of 22007 individuals, were recorded for the entire RDB.

**Figure 1.**
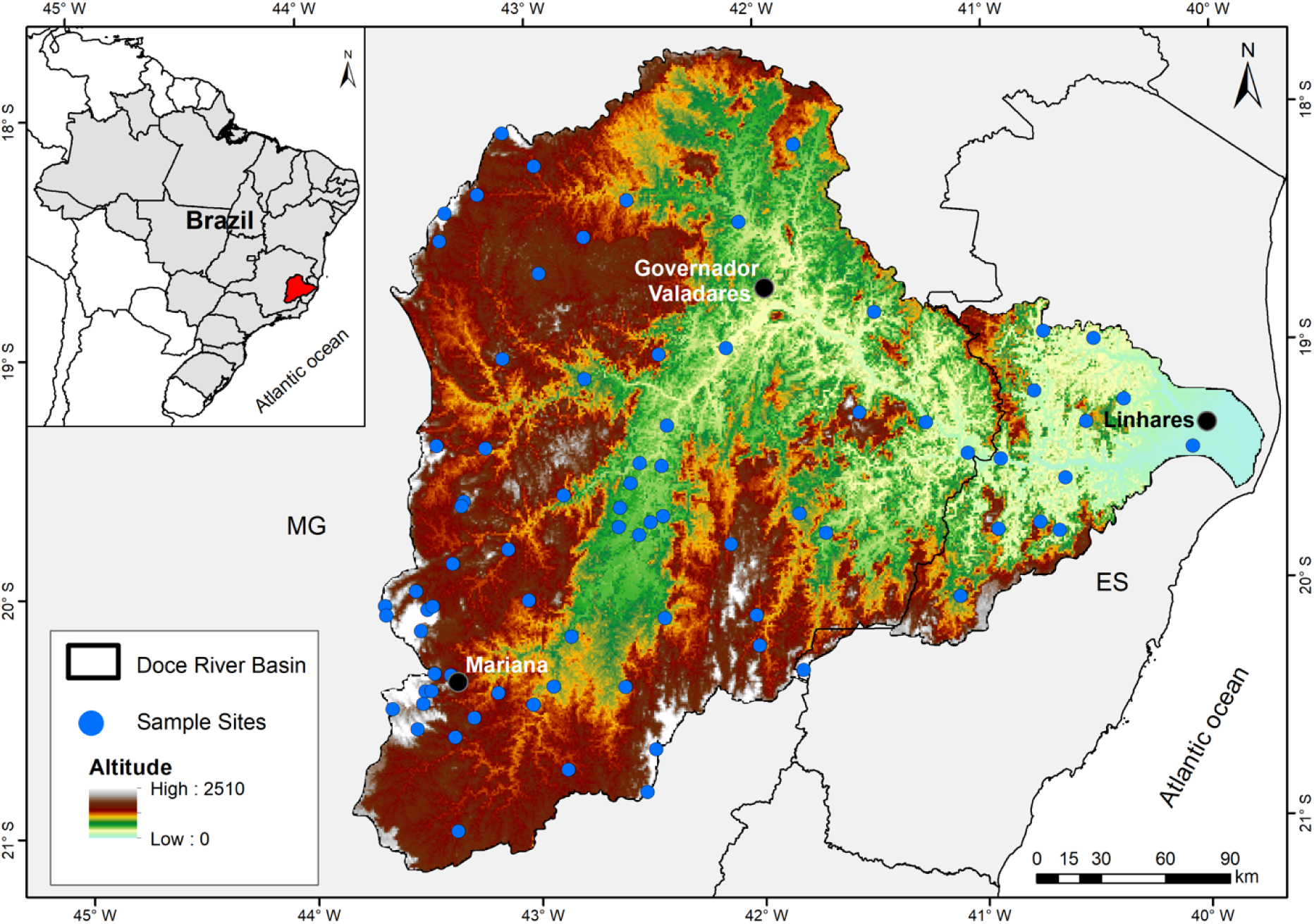
Rio Doce Basin (RDB) with the 78 studied forest sites (blue circle); three of the main cities within RDB (black circles). In the top left corner, the RDB (red) location in South America.

### Land-cover map and connectivity analysis

In this study, we used the classification of land use and occupation (see Fig 2A). The classification included 12 classes: native vegetation, pastures, reforestation area, rocky outcrops, open areas, agricultural areas, beaches, mining areas, urban areas, airports, roads and water. The RDB’s native vegetation has been largely suppressed by anthropogenic activity and most of the fragments are restricted to the steepest areas. In addition, pastures are also degraded with low soil coverage, soil compaction and intense trampling. (Agência Nacional de Águas, 2013).

**Figure 2.**
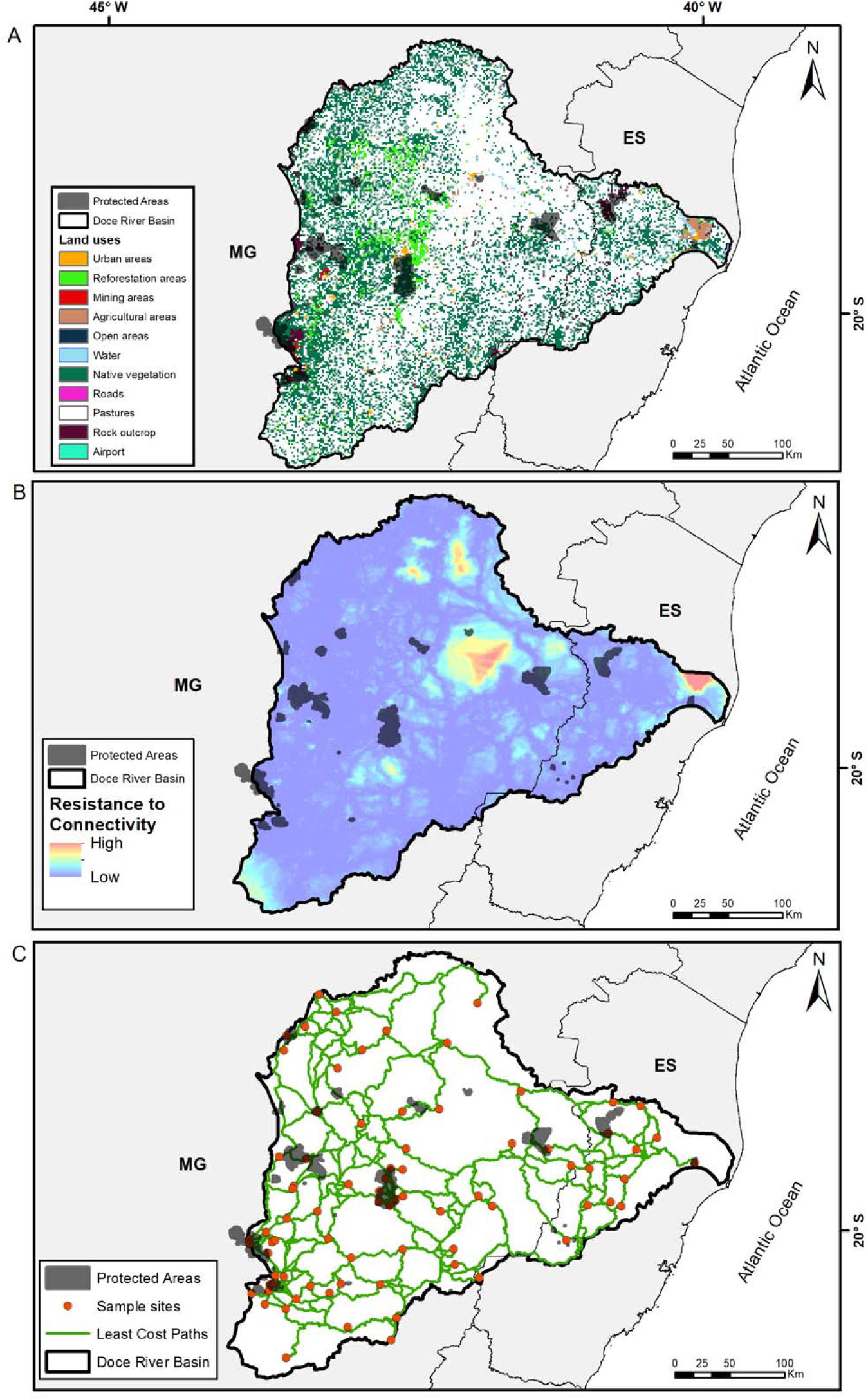
Maps generated based on resistance distance. (A) Land uses of the Rio Doce Basin provided by Agência Nacional de Águas. (B) Resistance to connectivity map based on the reclassification of land uses. (C) Least-Cost paths map. Black circles are the main municipalities, and red circles are the sample sites.

Our methodology was based on circuit theory (McRae and Kavanagh, 2006) We used the land cover map to build 6 hypothetical resistance surface models, where the elements on the land cover map were signed with a different resistance value, in the Linkage Mapper plugin from ArcGIS software (McRae et al., 2014). Therefore, each hypothetical resistance model has a different set of resistance values (Table 1. Appendices). The value of resistance added to a cell can be understood as the cost of moving through the cells (i.e., risk of mortality or difficulty). Native fragments are areas of low resistance and have been signed with a value of 1, while elements on the ground cover with higher resistance values, such as open areas, pastures and urban areas, have been signed with values greater than 50, with 100 as the maximum value.

**Table 1:** Models of Resistance created to model connectivity

| Resistance Value | Euclidean Distance | Model 1 | Model 2 | Model 3 | Model 4 | Model 5 |
| --- | --- | --- | --- | --- | --- | --- |
| 1 | - | Native Vegetation | Native Vegetation | Native Vegetation | Native Vegetation | Native Vegetation |
| 60 | - | Open Areas, Reforestation, Pastures | Reforestation, Rock Outcrops, Pastures |  | Open Areas, Reforestation, Rock Outcrops, Pastures | Open Areas, Rock Outcrops, Pastures |
| 70 | - | Rock Outcrop | Open Areas | - | - | Reforestation |
| 90 | - | Agriculture areas, beaches, mining areas, urban areas, airports, roads, | Agriculture areas, beaches, mining areas, urban areas, airports, roads, | - | Agriculture areas, beaches, mining areas, urban areas, airports, roads, | Agriculture areas, beaches, mining areas, urban areas, airports, roads, |
| 100 | - | Ocean, Water | Ocean, Water | All other classifications | Ocean, Water | Ocean, Water |

We used the 6 models as an input in Linkage Mapper Tool (McRae et al., 2014) to model connections between areas based on resistance and areas of central habitat (the 78 study sites sampled). The link mapper calculates the cost-weighted distance from one main area to another and these values are used to generate lower cost corridors and lower cost paths (LCP). The cost-weighted distance is normalized by the LCP between the areas, producing a normalized value of the lowest cost corridor so that the cells along the LCP are equal to 1. Then, the lowest cost normalized corridors are combined to generate the network of cost-distance corridors.

We inserted the protected areas (PA) within RDB to verify the effectiveness and relation of PA with the connectivity between fragments on the landscape scale.

### Data Analysis

We calculated a Jaccard similarity index matrix using the species composition of the studied forests (Figure 1). Then, we used the Jaccard index as a connectivity proxy between patches to assess the best resistance to promote connectivity. We created Generalized Linearized Models - GLM to test the relationship between the Jaccard index in pairs with the Euclidean Distance in pairs and the Lowest Cost Distance in pairs obtained from each of the 6 resistances to connectivity (Thiele et al., 2018a). We used quasi-binomial distribution, since the binomial models were overdispersed. We selected the best model in quasi-AIC values, where models with Δ quasiAIC> 2 were considered equally explanatory (Burnham et al., 2010). All models with ΔAIC less than 2 can be considered equally explanatory, so we used model 3 in our representation because it was based on a simpler resistance surface and because straight lines were not always possible in the landscape (Burnham and Anderson, 2002).

The analysis were done in R environment 3.4.3 version (R Development Core Team, 2017), where the Jaccard index was calculated with ‘vegdist’ function in ‘vegan’ package (Oksanen, 2016), the linearity of the models were tested with ‘cumres’ function in ‘gof’ package (González-Estrada and Villaseñor, 2018), the best model was assessed with ‘dredge’ function constraining the selection to models with only one predictive variable in ‘MuMIn’ package (Bartón, 2018), and the stats value of the models were tested with ‘Anova’ function in ‘car’ package, using the argument type = II (Fox et al., 2019).

## Results

In the selection of models, we found three models with quasi-ΔAIC <2. First, the model using Euclidean distance (P = 2.418e-06, AIC = 0.00). Second, model 3 in which we consider the forest to be the least resistant and all other types of land cover showed maximum resistance (p = 6.614e-07, quasi-ΔAIC = 1.59). Third, model 5, in which forests fragment when considered minimal resistance; Open areas, rocky outcrops, pastures and reforestation with intermediate resistance; and Agriculture areas, beaches, mining areas, urban areas, airports, roads, the ocean and water with high resistance values (p = 7.221e-07, quasi-ΔAIC = 1.61). All models with ΔAIC less than 2 can be considered equally explanatory, so we use model 3 in our representation because it was based on a simpler resistance surface and because straight lines are not always possible in the landscape. The map of RDB fragments shows that the hydrographic basin is highly fragmented, especially the areas close to the city of Governador Valadares and neighboring regions to the east, north and south (Fig. 2A). Most of the fragments are located in the west portion of RDB and promote greater connectivity for tree species in the RDB (Fig. 2C). The set of least-cost paths provide a very effective net of corridors between protected areas (Fig. 2C).

The regions that showed high levels of fragmentation, and less fragments showed high resistance to connectivity for tree species, especially the regions of Governador Valadares and, Linhares, and the extreme southwest of the RDB (Fig. 2B). Therefore, based on the map with lower cost corridors, these areas in the RDB have poor functional connectivity (Fig. 2C).

The result of the map of least cost paths shows that the areas around the municipality of Governador Valadares are not connected due to the greater resistance of the matrix, resulting from anthropogenic activity (mainly pastures and agricultural areas) that caused a reduction in fragments and, therefore, greater resistance of the landscape to connectivity.

## Discussion

Among the models we tested, the habitat/non-habitat model worked well in large areas due to its simplicity to apply and allowed to identify corridors and barriers across the landscape in the RDB. This model showed a main area with low connectivity for tree species resulting from intensive land use in the Governador Valadares region. Other regions also showed large areas with medium to high resistance to connectivity at the extreme east and extreme southwest RDB. For the most of the tree species, intensive land use results in hostile environments commonly found in agricultural landscapes around the world, such as arable fields and pastures, while areas suitable for dispersal and migration are rare (Thiele et al., 2018b). In the Governador Valadares region, much of the forest has been converted to pasture. The extreme east RDB and the extreme southwest RDB has been converted mainly into agricultural areas, *Eucalyptus* plantations and pastures. Therefore, these regions have gone through habitat loss, which increases the distance between patches, and increases the landscape resistance to connectivity (Hanski, 1999).

Patches of native vegetation in the RDB are abundant mainly in the west to northwest regions, restricted to steeper areas (Agência Nacional de Águas, 2013) due to mountainous topography in this region. Despite, the majority of the fragments have reduced size, small patches can be important for dispersers as stepping stones to reach bigger patches (Ribeiro et al., 2009). Furthermore, in the western portion of the RDB there is greater probability of connectivity (or less resistance for connectivity). Previous studies have shown the importance of small fragments or free-standing trees in the matrix for the maintenance of landscape connectivity (Luck and Daily, 2003; Matos et al., 2017; Mueller et al., 2014). On the other hand, the susceptibility to habitat loss and fragmentation is higher for species with higher demand of interior habitat and limited moving capacity (Laurance, 1990; Pfeifer et al., 2017).

Current plant distribution may not be a result of current landscape connectivity. For instance, the current diversity of plant species in grasslands in Sweden is not only a result of the current connectivity but it is related to the historical landscape connectivity (Lindborg and Eriksson, 2004). The patterns of plant dispersion in the past, when the basin was composed almost exclusively of forests, are rather an explanation of our model based on Euclidean distance. There can be a long time lag between landscape changes and the populations demise (Eriksson and Ehrlén, 2001) and it reflects the capacity of persistence of plant populations in isolated or in deteriorating environments (Lindborg and Eriksson, 2004). Thus, it is important to note that the current distribution of RDB plant species is also the result of the original distribution of native vegetation, historical occupation of the basin and dynamics of plant populations that respond to fragmentation.

The matrix surrounding fragments may affect the direction of dispersal. Certain land uses such as roads, waterways and agricultural areas act as barriers to some disperser preventing them to use these areas reducing the probability of dispersion (Taylor et al., 2006). For instance, northwards of Governador Valadares, there is an area that presents one connectivity path but there is high resistance to be connected southwards. As a consequence, important processes such as genes flow may be compromised, and it should be a priority area to manage because of that. Another important example is the extreme southwest RDB, where there is little connectivity in an area that has many springs of the Piranga River, the most important river among those that create the Doce River.

Landscape connectivity has three main components: 1) patterns and behaviors of species movement; 2) structure of resource patches (size and arrangement) and 3) the matrix in between patches (Taylor et al., 2006). The first two components are not always possible to be managed properly, since the inherent behavior of the species cannot be changed and the addition of areas to the fragments is often not viable due to political, social and economic constraints. The matrix is commonly greater in area than remnants of native vegetation. Thus, managing the matrix may be more important than managing only fragments in order to promote functional connectivity (Taylor et al., 2006). This may be the only remaining approach in the Governador Valadares region, in the extreme east and in the extreme southwest regions to increase habitat area and promote functional connectivity. Therefore, attention must be given to these regions in order to restore connectivity. Besides the Governador Valadares region has crucial importance, the extreme east region has fragments of Atlantic Forests with very high biodiversity, endemism and a high number of endangered species (Magnago et al., 2015; Matos et al., 2017). The extreme southwest of the basin presented relatively high level of resistance and one of the greatest areas without detectable least-cost paths in an area with high precipitation, supplying the basin with lots of water.

Functional connectivity for plants involves not only dispersal from source area, but also successful establishment and development in the receptor fragment (Auffret et al., 2017). Paths with enough area to receive propagules and restore the functional connectivity are crucial. A path that is well connected, in turn, will receive more dispersers and will provide also more propagules (Taylor et al., 1993). On one hand, regions with high resistance to connectivity and with few least-cost paths deserve a lot of attention in order to create least-cost paths suitable for tree species. On the other hand, the most important net of least-cost paths maintaining the connectivity between RDB’s fragments should be the priority for actions to maintain ecosystems functioning in the west to northwest portion. The actions should include prevention of clearcuttings and creation of reserves of ecosystems services that can be much cheaper, and effective than planting, and maintenance of corridors (Meira et al., 2016; Meira-Neto and Neri, 2017). Moreover, the west to northwest least-cost paths connect the most of the protected areas in the RDB. The connectivity is vital for protected areas, especially because Priority Areas for Biodiversity Conservation in the RDB accounts for 2,450,000 hectares (or 28% of RDB area) of which only 109,000 ha (4.45% of total Priority Areas) are in Integral Protection Conservation Units (Agência Nacional de Águas, 2013). This is a very low amount of reserves, and new well-connected reserves of ecosystem services should be created.

## Conclusions

The results of modeling landscape connectivity using empirical data have proven to be an effective tool for prioritizing sites of greater connectivity in fragmented landscapes. Despite the highly fragmented landscape found in the RDB, our approach allows to highlight that there is still a relatively well-connected west-northwest region and functioning to ensure the persistence of plant species of tree species. Nevertheless, the central region of Governador Valadares, the neighboring regions, the regions of the extreme east and southwest of the RDB were strongly affected by habitat loss, have great resistance to connectivity, and few least cost paths. Therefore, land restoration and reclamation projects in these areas should be encouraged to mitigate the lack of connectivity, ensuring the persistence of plant populations, maintaining biodiversity and vital processes for ecosystems. Thus, we defend that public policies should focus on the restoration of degraded areas on landscapes with high resistance to connectivity, while reinforce conservation actions of less degraded areas to preserve functioning and connectivity in the RDB.

## 7. Appendix

